# A Comprehensive Benchmark Study on Biomedical Text Generation and Mining with ChatGPT

**DOI:** 10.1101/2023.04.19.537463

**Authors:** Qijie Chen, Haotong Sun, Haoyang Liu, Yinghui Jiang, Ting Ran, Xurui Jin, Xianglu Xiao, Zhimin Lin, Zhangming Niu, Hongming Chen

## Abstract

In recent years, the development of natural language process (NLP) technologies and deep learning hardware has led to significant improvement in large language models(LLMs). The ChatGPT, the state-of-the-art LLM built on GPT-3.5, shows excellent capabilities in general language understanding and reasoning. Researchers also tested the GPTs on a variety of NLP related tasks and benchmarks and got excellent results. To evaluate the performance of ChatGPT on biomedical related tasks, this paper presents a comprehensive benchmark study on the use of ChatGPT for biomedical corpus, including article abstracts, clinical trials description, biomedical questions and so on. Through a series of experiments, we demonstrated the effectiveness and versatility of Chat-GPT in biomedical text understanding, reasoning and generation.

## 1 Introduction

In recent years, there has been a tremendous growth in the field of natural language processing (NLP) and machine learning. One of the most significant advancements in NLP is the development of large language models such as GPT (Generative Pre-trained Transformer)(Radford et al., 2018, 2019; Brown et al., 2020) and its various variants, which have shown remarkable performance in a number of language tasks. Usually GPT models were initially pre-trained on massive text data and then fine-tuned on specific downstream tasks to generate human-like languages.

In the domain of biomedical text mining, NLP techniques have demonstrated the potential to revolutionize research and clinical practice. However, the complexity of biomedical language and the vast amount of data size still make it a challenging task to develop robust models for text generation and mining. In this paper, we present a comprehensive benchmark study on evaluating the performance of ChatGPT model(Ouyang et al., 2022), a large-scale GPT-based language model, for biomedical text generation and mining.

The paper is organized as following. Firstly, we will provide an overview of the related work in biomedical textmining and highlight the strength and limitations of current approaches. Secondly, the ChatGPT model and its applications in NLP will be described. Thirdly, we will discuss the benchmarking and experimental protocols conducted in this study. Finally, we will present the performance of ChatGPT in various biomedical text generation and mining tasks along with other baseline biomedical NLP models and discuss the potential applications and future directions of ChatGPT in biomedical research and clinical practice.

Overall, this paper aims to contribute to the growing body of research in the field of biomedical NLP by providing a comprehensive evaluation of ChatGPT model on biomedical text generation and mining. By comparing the performance of ChatGPT with other SOTA biomedical models on several biomedical related NLP benchmark sets, we hope to provide the pros and cons of ChatGPT model in dealing with biomedical related tasks, which may inspire further development of more advanced NLP models for biomedical data analysis.

## 2 Background and Related Work

In recent years, natural language processing (NLP) techniques have gained significant attention in the biomedical domain due to the vast amount of textual data generated by scientific publications, electronic health records, and social medias etc. Biomedical text mining, a sub-field of NLP, aims to extract, analyze and summarize useful information, and derive insightful knowledge from either structured or unstructured biomedical texts. Usually, extracting knowledge from biomedical text requires substantial human effort and is time-consuming. Thus, automated text generation and mining techniques can greatly assist researchers via extracting or deriving valuable insights from the available big data in biomedical literature.

Recently, one of the most promising advances in NLP field is the development of so called largescale language models (LLMs) to using hundreds of billions of parameters and to training on gigabytes of text (Brown et al., 2020; Ouyang et al., 2022). These models have been shown to achieve state-of-the-art (SOTA) performance in several NLP tasks, including text generation, question and answering (QA), and text summarisation. The capability of these models to generate coherent and contextually relevant text makes them ideal candidates for biomedical text generation and mining. By identifying critical data points for clinical trials and drug discovery, LLMs can assist in advancing the creation of new drugs and treatment approaches.

Several studies have demonstrated the potential of these language models in biomedical text mining. For instance, BioLinkBERT was an LM pretraining method that leverages links between biomedical documents. SciFive (Phan et al., 2021) applied a domain-specific T5 model (Raffel et al., 2020) that has been pre-trained on large biomedical corpora.

Moreover, *pre-train, prompt and predict* (Liu et al., 2023) is an emerging paradigm for applying LLMs to new problems without fine-tuning the weights on the task. Prompt-based learning involves enhancing the problem statement with specific instructions so that the model’s response to the prompt results in a solution. This methodology enables LLMs to learn from a limited set of examples, referred to as *shots*, which are integrated into the prompts themselves(Brown et al., 2020). ChatGPT (Ouyang et al., 2022) has garnered enormous attention due to its remarkable success in instruction understanding and human-like response generation. According to recent research, the Chat-GPT language model created by OpenAI has shown promising results in performing at par with humans on MBA exams conducted by the Wharton Business School.(Rosenblatt, 2023)This indicates that AI language models like ChatGPT have the potential to compete with human knowledge and could be utilized to assist professionals.(Choi et al., 2023; Baidoo-Anu and Owusu Ansah, 2023). Also, their impressive performance on diverse NLP tasks, coupled with their ability to generalize to unfamiliar tasks, highlights their potential as a versatile solution for a variety of challenges in natural language understanding, text generation, and conversational AI.

While these studies have demonstrated the potential of LLMs in biomedical text mining, there is still a lack of comprehensive evaluation of LLMs on broad biomedical tasks. This study aims to provide a large scale study of the latest ChatGPT model in biomedical text generation and mining. We investigated the performance of ChatGPT in several biomedical NLP tasks, including entity recognition, paragraph summarization, and answer generation etc. We also explored the possibility of using ChatGPT to assist researchers in extracting useful knowledge from the available biomedical data.

Further, (Wei et al., 2022) demonstrates that LLMs could be achieved by generating a *chain of thought*(a series of intermediate reasoning steps) to improve the ability of large language models to perform complex reasoning, coined “*Chain-of-Thought*” (CoT). This prompt not only appears to expose valid reasoning but also translates into superior zero-shot performances. (See example in **Results and Discussions**.)

## 3 ChatGPT for Biomedical NLP

The volume of biomedical literature has significantly expanded in recent years, leading to a urging need for robust text mining tools for biomedical application. Numerous studies have shown that pre-trained language model can help accelerate the progress of general biomedical NLP applications.

A common workflow for training domain specific language model is to pretrain models on large general data sets to learn general features of languages and then fine-tune on more focused domain specific data. Large models, e.g. BERT-based or GPT-based models, were firstly pretrained with huge amount of text data either supervisedly, semisupervisedly or unsupervisedly. The pre-trained models offer representation, or in another word, featurization for the input text, which is regarded as general understanding of the model for the input sentences. Then for any downstream task, the pre-trained model is combined with a prediction head and fine-tuned together with a relatively small domain specific train set in a supervised pattern. In some studies, the parameters of the pre-trained model may also be frozen. The prediction head gives a desired output that can be utilized to evaluate the model performance. ChatGPT is a generative model based on GPT-3.5 and fine-tuned to accomplish text generating tasks. As the exact model structure and parameters are not released by OpenAI yet, it is impossible to directly fine-tune the model toward user supplied data. However, it has been shown that ChatGPT can achieve human-like dialogue results through chatting with specifically engineered prompts. Here, we employed prompt engineering method to engage ChatGPT model in biomedical related NLP tasks and then evaluate its performance. In most of cases, the ChatGPT model was challenged in a zero-shot or few-shot manner (as part of the prompt).

The design of the prompt is crucial for the output of ChatGPT. In general, the prompt should at least consist of a body of background context, an instruction part telling ChatGPT what’s the task supposed to be done, and a constrain part for formating the output and content. For instance in a yes/no QA task, ChatGPT should be told to ‘answer in a simple yes or no’ so that we can obtain structured results and calculate performance metrics. But there are still cases that output of ChatGPT does not obey the constrains, e.g. supplying reasons after a ‘yes’ for a QA task or answering entities that does not exist in the text for a named entity recognization (NER) task. To deal with these exceptions, we choose to judge the answer at first and then emphasize again the constrains. This requires multiple rounds of question and answering. Figure 1 provides An example of zero-shot biomedical NLP task using ChatGPT.

**Figure 1:**
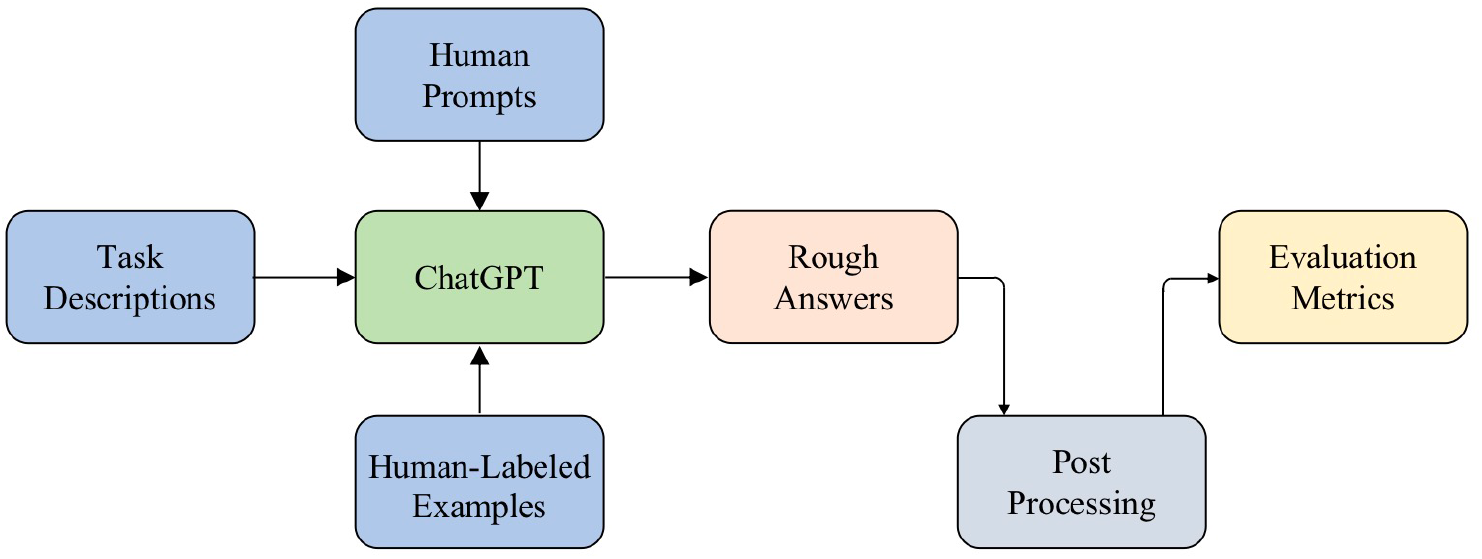
An overview of the workflow for Biomedical NLP using ChatGPT.

## 4 Experiments

We applied the pattern proposed in Section 3 to test the performance of ChatGPT on Biomedical NLP tasks. Considering the model accessibility and computation speed, we tested the ChatGPT model built on GPT-3.5 to evaluate the performance on Biomedical NLP tasks. In this section, we will first introduce the benchmark data sets, followed by a description of our evaluation tasks and their respective implementation details. Finally, we will present the results of ChatGPT.

### 4.1 BLURB Benchmark

#### BLURB

We utilized a comprehensive benchmark data set for Biomedical NLP, the *Biomedical Language Understanding & Reasoning Benchmark (BLURB)*^1^, which is an extensive collection of biomedical NLP tasks derived from publicly accessible data sources and contains 13 biomedical NLP subsets grouped in six types of task. These tasks include NER, evidence-based medical information extraction (PICO), biomeidical relation extraction(BRE), sentence similarity, document classification, and QA. An overview of the BLURB datasets can be found in Table 1.

**Table 1:** Overview of the BLURB benchmark. We list the numbers of instances in train, dev, and test, as well as their respective evaluation metrics.

| Dataset | Task | Train | Dev | Test | Evaluation Metrics |
| --- | --- | --- | --- | --- | --- |
| BC5-chem | NER | 5203 | 5347 | 5385 | F1 entity-level |
| BC5-disease | NER | 4182 | 4244 | 4424 | F1 entity-level |
| NCBI-disease | NER | 5134 | 787 | 960 | F1 entity-level |
| BC2GM | NER | 15197 | 3061 | 6325 | F1 entity-level |
| JNLPBA | NER | 46750 | 4551 | 8662 | F1 entity-level |
| EMB PICO | PICO | 339167 | 85321 | 16364 | Macro F1 word-level |
| ChemProt | BRE | 18035 | 11268 | 15745 | Micro F1 |
| DDI | BRE | 25296 | 2496 | 5716 | Micro F1 |
| GAD | BRE | 4261 | 535 | 534 | Micro F1 |
| BIOSSES | Sentence Similarity | 64 | 16 | 20 | Pearson |
| HoC | Document Classification | 1295 | 186 | 371 | Acerage Micro F1 |
| PubMedQA | QA | 450 | 50 | 500 | Accuracy |
| BioASQ | QA | 670 | 75 | 140 | Accuracy |

**Table 2:**
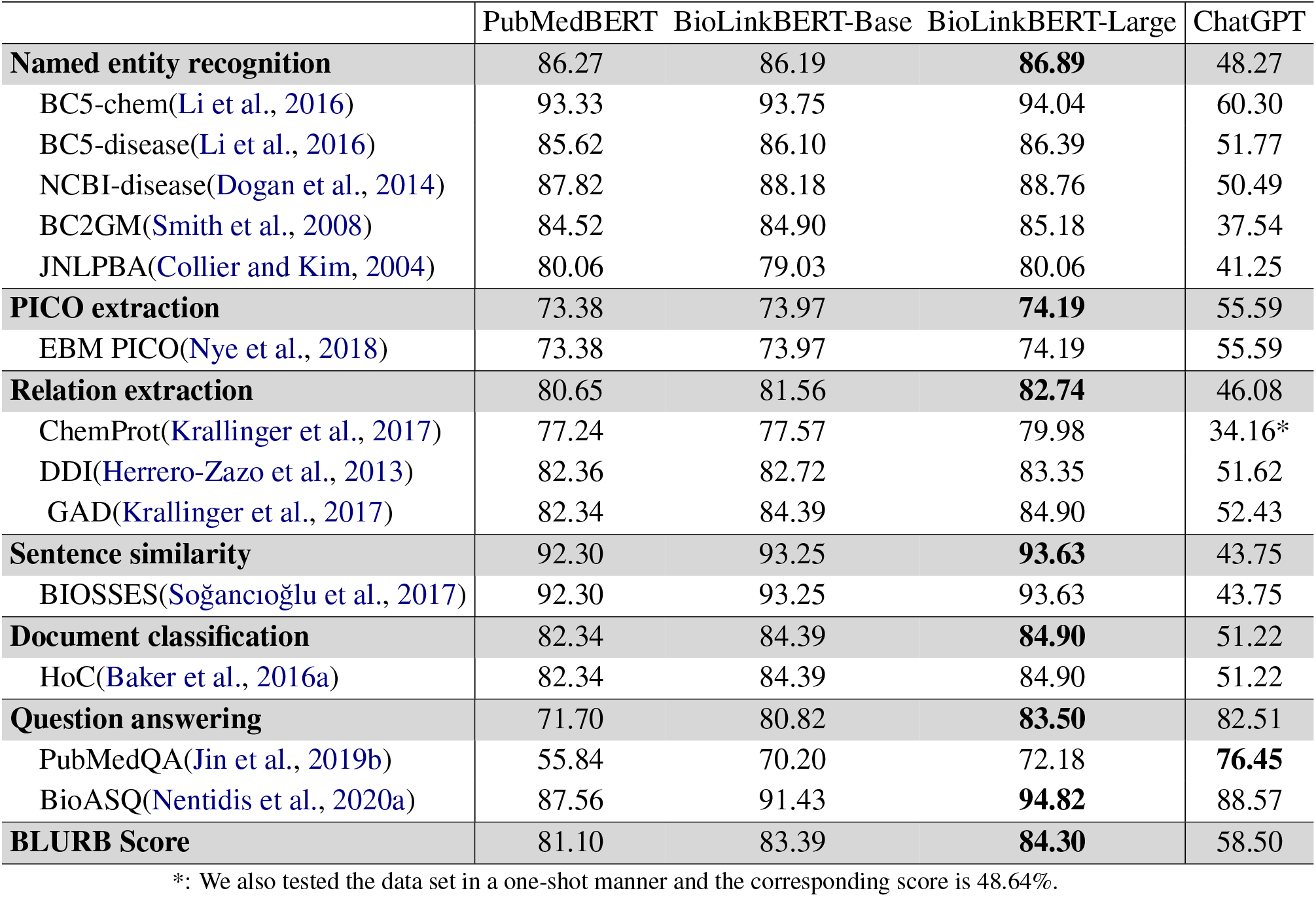
Performance on BLURB benchmark. We list the overall BLURB Score and the score for each task in gray shaded cells.

#### Evaluation Metrics

To calculate the overall score for BLURB, the simplest approach would be to report the average score across all tasks. However, this may be biased by some high-scored tasks. Therefore, we provided both average score per task class which reflect the performance on data sets belonging to the same task type, and the average overall score among all task types.

### 4.2 Biomedical NLP Tasks

In order to achieve optimal performance for ChatGPT model across different tasks, specific prompts for various tasks were designed based on the pattern proposed in Section 3.

#### 4.2.1 Named Entity Recognition

NER task is a process for identifying and predicting named entities, such as name of chemical substance, disease, gene, and protein, within given input text. Five NER datasets from the BLURB benchmark were investigated, including *BC5-Chemical, BC5-Disease, NCBI-Disease, BC2GM*, and *JNLPBA*. For these datasets, the same splits for train, validation, and test set as utilized by (Crichton et al., 2017) were used in current study. BC2GM is a corpus data set, which consists of over 20,000 abstracts and full-text articles from the MEDLINE database published during the period 1991-2003. Each document in the corpus was annotated by domain experts with gene names and synonyms, as well as their corresponding Entrez Gene IDs. The NER task on the BC2GM dataset requires a predictive model to identify all gene entities mentioned in a text (Smith et al., 2008). The BC5-chem and BC5-disease data sets were retrieved from the BioCreative challenge and were respectively designed for NER tasks towards chemical and disease entities. The former data set contains over 1,500 documents with approximately 42,000 chemical annotations, while the latter one contains over 1,500 documents with approximately 24,000 disease annotations. The NCBI-disease corpus was created by the National Center for Biotechnology Information (NCBI) for disease recognition tasks in biomedical natural language processing (Doğan et al., 2014). It consists of over 20,000 PubMed abstracts that were manually annotated by domain experts with disease names and their corresponding disease IDs from the Medical Subject Headings (MeSH) vocabulary. The JNLPBA (Joint Workshop on Natural Language Processing in Biomedicine and its Applications) corpus was provided by the JNLPBA conference specifically for gene entity recognition (Collier and Kim, 2004). It consists of over 2,000 PubMed abstracts, manually annotated by domain experts. These corpora cover a diverse range of biomedical topics, making it a valuable resource for training and evaluating machine learning models for NER tasks. In the BLURB, the annotation format in the corpus was unified for five NER data sets. Specifically, a pair of entity type masks were added before and after the words representing the entity name. For example, the mask “gene* [entity] *gene” was inserted to the text to label the gene entity in the bracket. The disease and chemical entities were masked similarly. In this study, Chat-GPT was employed to recognize the entity name in the text without any prior knowledge. The prompt was designed as:

*Paragraph: <Paragraph ID>* | *<text> Please extract all chemicals/genes/diseases mentioned in the paragraph. Answer with the format “<Paragraph ID>* | *<recognized entities>“*

#### 4.2.2 PICO

PICO stands for Patient/Population, Intervention, Comparison and Outcomes. PICO model is used to construct a clinical question. The practice of evidence based medicine (EBM) aspires to inform healthcare decision using the total relevant evidence.(Nye et al., 2018) *EBM-NLP* is an biomedical corpus comprising 4993 medical abstracts describing clinical trials, containing spans of token corresponding to three categories, ie. Populations, Interventions and Outcomes in the clinical trial. Each P/I/O span is further annotated with more detailed labels, e.g. Age, Sex information etc.(Huang et al., 2006). The test set contains 191 abstracts where 16364 out of around 54000 tokens are related to P/I/O categories and others are labeled as “None”. Comparison(C) is not annotated in this corpus. This is like a token-wise multi-classification task as typical classifiers did. But it is inconvenient to ask ChatGPT to classify each word one by one. In practice, we designed prompts similar to the NER tasks for asking ChatGPT to extract all the words related to P/I/O class and the rest of words were attributed as ‘None’. A natural-language-like prompt was designed as:

*Reference: <abstract> The reference describe a clinical trial. Which words are about the participants/interventions/outcomes? You can only answer with words or phrase in the reference. If nothing mentioned, answer “None”*.

The PICO task is somehow similar to a NER task but there are still some differences between the tasks. For example, the words annotated as P/I/O can not only be entity names, but also sentences composed of prepositions, adverbs and even punctuations etc, which describe the target span. As the result is evaluated with macro word-level F1 score, a neural network classifier can make a prediction for each token(word), but it’s impractical for ChatGPT, a generative model, to do the task in such a word-wise way. For example, ChatGPT only answers a word for one time even if the word appears several times in the abstract. In order to properly evaluate the performance of ChatGPT, these words were weighted with the number of appearance when counting the confusion matrix, and the punctuations were excluded.

#### 4.2.3 Biomedical Relation Extraction

Biomedical relation extraction (BRE) task focuses on identifying and extracting relationships between medical entities in input text, such as connections between diseases and drugs, or symptoms and treatments. Formally, let *x* represents a sentence containing two medical entities, *e*_1_ and *e*_2_, with *r* being the relation between them. The BRE task can be framed as a classification problem, where the objective is to learn a function *f* (*x, e*_1_, *e*_2_) *→ r*, with *r* belonging to the set of possible relations *R*. This function leverages the context provided by sentence *x* to predict the relation between entities (*e*_1_, *e*_2_). The performance of BRE models is generally assessed using standard classification metrics, such as confusion matrix based Precision, Recall, and F1-score etc.

We evaluated the performance of ChatGPT on three biomedical datasets: ChemProt, DDI, and GAD.

To assess ChatGPT’s ability in the BRE task, for example, we crafted a prompt (for the GAD Dataset) as following: *“Does the reference indicate a relationship between the @DISEASE$ and the @GENE$ without specifying the exact disease and gene? Response with “yes” or “no”*.*”* By doing in this way, it allowed us to gauge ChatGPT’s effectiveness in recognizing and extracting relationships between medical entities within the context of biomedical text.

As indicated by its name, ChemProt is a data set containing around 700000 unique chemicals, 3000 proteins and 2,000,000 interactions overall from around 2500 documents. All the interactions are grouped into 10 groups according to biological semantic classes. A five-group-subset was used as the test set. The five groups in the test set include: 1) upregulator | activator | indirect upregulator, 2) downregulator | inhibitor | indirect downregulator, 3) agonist | agonist-activator | agonist-inhibitor, 4) antagonist, 5) substrate | product of | substrate product of. There are even more groups in the train set and validation set. Besides, unrelated chemical substance and protein pairs were labeled as ‘None’ to enrich the data set. Clearly, domain knowledge is required to help understand what the exact relation means and might be missing in general LLMs like ChatGPT. To overcome the difficulty, we test ChatGPT by adding one sample of the validation set for each group into the prompt, in another word, with the one-shot learning manner.

The Drug-Drug Interaction corpus(Herrero-Zazo et al., 2013) was created to facilitate research on pharmaceutical information extraction, with a particular focus on pharmacovigilance. It contains sentence-level annotation of drug-drug interactions on PubMed abstracts.

Gene-disease associations database (GAD) (Bravo et al., 2015)set is a collection of around 5000 published gene/disease associations. The gene name and disease name in the document are recognized and masked. Here, the label indicates whether the document implies an association between the gene and the disease as a binary classification task. Different from ChemProt, the relations were not strictly defined with a biology terminology and could be ambiguous sometimes. 534 sentences were used as the test set.

#### 4.2.4 Sentence Similarity

The Sentence Similarity task involves predicting a similarity score based on the likeness of a given pair of sentences. The BLURB benchmark contains the *BIOSSES* dataset consisting of 100 pairs of sentence from Text Analysis Conference(TAC) Biomedical Summarization Track (Soğancioğlu et al., 2017). The train, validation, and test splits were the same with the ones used before (Peng et al., 2019) and we tested ChatGPT on a test set of 20 pairs. The score is in the range of 0-5. The definition is declared in the table 3. Each sample was scored by 5 annotators and the average score was used as the ground truth, leading to a regression-like task. The prompt is designed as: *What is the similarity score between the <sentence1> and the <sentence2>? Response with float ranging from 0 (no relation) to 4 (equivalent)?*

**Table 3:**
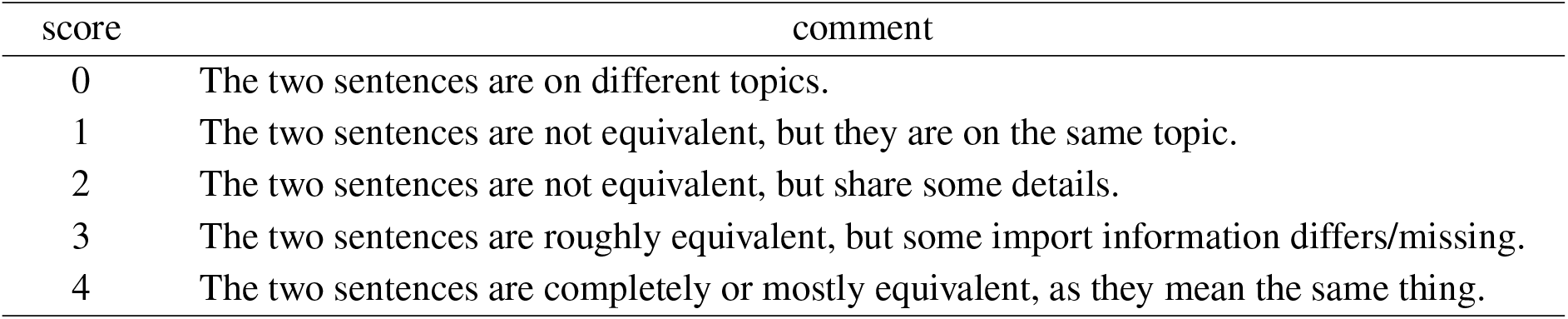
Definition of the scores in the BIOSSES data set

**Table 4:** Metrics for five NER tasks with BLURB benchmark datasets

| NER Task | F1-score | Recall | Precision |
| --- | --- | --- | --- |
| BC5-disease | 0.52 | 0.59 | 0.46 |
| BC2GM | 0.38 | 0.46 | 0.32 |
| BC5-chem | 0.60 | 0.76 | 0.50 |
| NCBI-disease | 0.50 | 0.51 | 0.50 |
| JNLPBA | 0.41 | 0.55 | 0.33 |

**Table 5:**
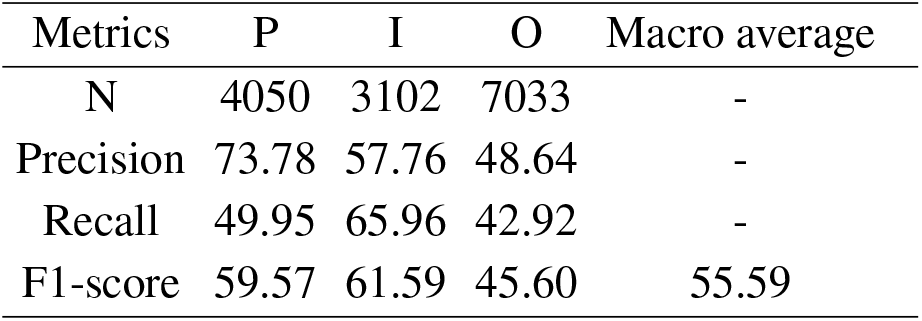
Performance of ChatGPT on EBM PICO task. Annotated punctions are excluded

#### 4.2.5 Document Classification

Document Classification is a procedure of assigning one or more pre-defined labels to a document. Evaluation for this task was done at the document level, ie. aggregating labels across all sentences within a document. We utilized the HoC data set from the BLURB benchmark, which was curated by (Baker et al., 2016b) and employed the same splits of train, validation, and test set.

We have designed the following prompt to enable ChatGPT to carry out the document classification task: “*document: <text>; target: The correct category for this document is ? You must choose from the given list of answer categories (introduce what each category is* …).”

#### 4.2.6 Question Answering

The QA task refers to predicting answers under the given context, in which the first sentence is question. Answers are either two labels (yes/no) or three labels (yes/maybe/no). We utilized the PubMedQA (Jin et al., 2019a) and BioASQ (Nentidis et al., 2020b) data sets for evaluation. For both data sets, the original train, validation, and test splits within the BLURB benchmark were used.

For evaluation of ChatGPT on PubMedQA and BioAS, we simply designed the following prompt: *“question: <text>; context: <text>; answer: <text>; target: the answer to the question given the context is (yes or no)? “*

### 4.3 Results and Discussions

We tested the performance of ChatGPT with engineered prompts as mentioned in previous sections and altogether, six types of biomedical text mining task (NER, PICO, BRE, Sentence Similarity, Document Classification and QA) were explored.

### Baseline models

We selected three baseline models that are SOTA on the BLURB benchmark for comparison with ChatGPT, ie. **PubmedBERT, BioLinkBERT-Base** and **BioLinkBERT-Large**. All the models are based on the BERT architecture. **PubmedBERT**(Gu et al., 2021) was pre-trained on PubMed and **BioLinkBERT-Base**(Yasunaga et al., 2022) was pre-trained on PubMed with citation links. The **BioLinkBERT-Large** model was specifically pre-trained on a large corpus of biomedical literature and clinical notes, which allow to capture the complex terminology and domain-specific knowledge required for biomedical NLP tasks. It contains over 335 million parameters, making it one of the largest pre-trained models in the biomedical domain.

Table 2 shows the performance of ChatGPT and baseline models on BLURB benchmark. Although, in general, ChatGPT got a BLURB score of 59.46 which is significantly worse than the SOTA baselines, there are still interesting conclusions can be drawn for ChatGPT. On the other hand, we should bear in mind that ChatGPT was trained as a general language model, while the baselines are models particularly trained on biomedical corpus.

Among all types of task, QA task is the only type of task that ChatGPT is comparative to the baselines. In this case, ChatGPT (82.5) outperforms PubMedBERT (71.7) and BioLinkBERT-Base (80.8) and is very close to the BioLinkBERT-Large (83.5). In particular on the PubMedQA data set, ChatGPT exceeded the baselines significantly and the score is close to the human performance of 78.2% (Jin et al., 2019a) and the SOTA score of 79.6% (He et al., 2022). This results suggest that ChatGPT has strong capability in understanding these questions and is also able to give simple answers as good as human do.

The NER tasks in BLURB are to identify entities of chemical substance, disease and gene name. The recognition accuracy of ChatGPT among various data sets is, from high to low, chemicals (BC5-chem) > diseases (BC5-disease and NCBI-disease) > genes(BC2GM and JNLPBA), which is consistent with the baselines. This trend reflects that disease and gene name have higher intrinsic complexity than chemical name. We attributes the poor performance of ChatGPT to the missing of supervised training and lack of training data in biomedical field. As far as we know, ChatGPT was trained mainly on the data of web sites, social media posts, books and articles. But biomedical entities, especially terminologies, are uncommon in the daily usage. It’s probably explainable that ChatGPT does not understand well these texts which need more domain knowledge to interpret.

As introduced in the Section 4.2.2, PICO task is similar to NER. ChatGPT performs worse than the baselines but the gap is smaller comparing to NER tasks. PICO task was assumed to be easier since many of the target words/sentences are commonly used in daily life and easy to understand. One possible reason for the poor performance is that ChatGPT may miss the short sentences or phrases while can successfully extract the long ones. Among following phrases and sentences labeled as P class in a document ‘treated hypertensive patients’, ‘hypertensive patients receiving drug treatment’, ‘hypertensives on chronic, stable antihypertensive therapy’, ‘people with one or more cardiovascular risk factors’, ‘hypertensives under treatment’, ‘Fifteen Italian hypertension units studied 142 hypertensive patients(76 men, 66 women; mean age 59+/-5.9 years) treated with different antihypertensive drugs’, ChatGPT failed to recognize those short phrases/sentences as P class and only labeled the last long sentence correctly.

Relation extraction tasks require a model to be able to identify the relation of a pair of entities masked in the text. For DDI and GAD data sets, whose format is similar to QA tasks requiring the model output to be a simple ‘yes’ or ‘no’, ChatGPT performed poorly. ChemProt set is more complex due to the requirment of grouped relation and ChatGPT got even lower score than other tasks. A straightforward guess is that the so-called ‘relation’ is not that clear. It’s hard for ChatGPT to understand what the relation mentioned in our prompt refers to. To validate the guess, we tested ChatGPT on ChemProt in one-shot manner, in which one sample for each relation group was provided. The one-shot method greatly improved the score from 34.16% to 48.64%. Another thing we noticed from the results was that ChatGPT tended to be confused by other words in the text and often assigned relation labels to entity pairs which are actually unrelated. The original ChemProt data contains only 3458 test samples where the entity pairs are all related. While in BLURB benchmark set, this set was augmented with 15745 negative pairs. It was found that false positive rate(FPR) is as high as 75%. We tested ChatGPT on the original ChemProt data and the F1-score is 79.93%. Through these experiments, we expect that ChatGPT still has room to do a better job on these tasks with more carefully designed prompts, e.g. supplying more instructions about the relation that the data set concerns and add more shots.

The document classification task is quite challenge for ChatGPT. On one hand, the number of the answer category is uncertain, it may be an empty category, it may be one of the categories, or it may be multiple categories. On the other hand, this few shot learning scenario is not friendly for ChatGPT, as it is really difficult to understand the labels without enough domain knowledge. It can be seen from Table 2 that on the HoC data set, ChatGPT only obtained an F1 value of 51.22%, which is much worse than BERT based models, indicating that the performance of ChatGPT in processing medical text classification tasks with few samples is still far from optimal.

Sentence similarity is also a difficult case for ChatGPT with zero-shot. Different from other tasks, the similarity defined on the BIOSSES data is quite subjective and the similarity score could be ambiguous. The Y variable is the average score from 5 annotators and the human opinions are always diverse. The score deviation of a certain pair of sentences could be up to 2. The pearson coefficient between individual annotations and the ground truth is only 0.5. So, in this sense, ChatGPT performed actually not worse than human. The baselines got a high score due to the fine-tune process. ChatGPT may work better on this task if we fed some samples from the train set within the prompt. As we focused mainly on the zero-shot method and tried to evaluate the overall capacity of the ChatGPT, we did not test this strategy for this small data set with only 100 pairs of sentences.

## 5 Conclusion

Based on our experiments, the ChatGPT built on the early version of GPT-3.5 performed poorly on several biomedical NLP benchmark data sets. The biomedical domain is clearly a challenging professional field to deal with for a general LLM running in the zero or few shot scenario. Another common problem is that ChatGPT is a generative model while most benchmark sets are designed for supervised models, requiring a structured prediction. SOTA language models are usually fine-tuned in a supervised manner based on a pre-trained large model. Though we can add instructions in the prompt to constrain the output of the ChatGPT, there are still chances that the ChatGPT output doesn’t follow the expected format. Having said that, the superior version GPT-4 has recently been released and demonstrated better ability of natural language understanding and reasoning. We are looking forward to test newer version of ChatGPT on professional NLP tasks to explore the potentiality of LLMs.

## Footnotes

1 https://microsoft.github.io/BLURB/

## Notes

### Competing Interest Statement

The authors have declared no competing interest.

## References

David Baidoo-Anu and Leticia Owusu Ansah. 2023. Education in the era of generative artificial intelligence (ai): Understanding the potential benefits of chatgpt in promoting teaching and learning. Available at SSRN 4337484.

Simon Baker, Ilona Silins, Yufan Guo, Imran Ali, Johan Högberg, Ulla Stenius, and Anna Korhonen. 2016a. Automatic semantic classification of scientific literature according to the hallmarks of cancer. Bioinformatics, 32(3):432–440.

Simon Baker, Ilona Silins, Yufan Guo, Imran Ali, Johan Högberg, Ulla Stenius, and Anna Korhonen. 2016b. Automatic semantic classification of scientific literature according to the hallmarks of cancer. Bioinform., 32(3):432–440.

Àlex Bravo, Janet Piñero González, Núria Queralt-Rosinach, Michael Rautschka, and Laura Inés Furlong. 2015. Extraction of relations between genes and diseases from text and large-scale data analysis: implications for translational research. BMC Bioinform., 16:55:1–55:17.

Tom Brown, Benjamin Mann, Nick Ryder, Melanie Subbiah, Jared D Kaplan, Prafulla Dhariwal, Arvind Neelakantan, Pranav Shyam, Girish Sastry, Amanda Askell, et al. 2020. Language models are few-shot learners. Advances in neural information processing systems, 33:1877–1901.

Jonathan H Choi, Kristin E Hickman, Amy Monahan, and Daniel Schwarcz. 2023. Chatgpt goes to law school. Available at SSRN.

Nigel Collier and Jin-Dong Kim. 2004. Introduction to the bio-entity recognition task at jnlpba. In Proceedings of the International Joint Workshop on Natural Language Processing in Biomedicine and its Applications (NLPBA/BioNLP), pages 73–78.

Gamal K. O. Crichton, Sampo Pyysalo, Billy Chiu, and Anna Korhonen. 2017. A neural network multi-task learning approach to biomedical named entity recognition. BMC Bioinform., 18(1):368:1–368:14.

Rezarta Islamaj Dogan, Robert Leaman, and Zhiyong Lu. 2014. NCBI disease corpus: A resource for disease name recognition and concept normalization. J. Biomed. Informatics, 47:1–10.

Rezarta Islamaj Doğan, Robert Leaman, and Zhiyong Lu. 2014. Ncbi disease corpus: a resource for disease name recognition and concept normalization. Journal of biomedical informatics, 47:1–10.

Yu Gu, Robert Tinn, Hao Cheng, Michael Lucas, Naoto Usuyama, Xiaodong Liu, Tristan Naumann, Jianfeng Gao, and Hoifung Poon. 2021. Domain-specific language model pretraining for biomedical natural language processing. ACM Transactions on Computing for Healthcare (HEALTH), 3(1):1–23.

Pengcheng He, Baolin Peng, Liyang Lu, Song Wang, Jie Mei, Yang Liu, Ruochen Xu, Hany Hassan Awadalla, Yu Shi, Chenguang Zhu, Wayne Xiong, Michael Zeng, Jianfeng Gao, and Xuedong Huang. 2022. Zcode++: A pre-trained language model optimized for abstractive summarization.

María Herrero-Zazo, Isabel Segura-Bedmar, Paloma Martínez, and Thierry Declerck. 2013. The ddi corpus: An annotated corpus with pharmacological substances and drug–drug interactions. Journal of biomedical informatics, 46(5):914–920.

Xiaoli Huang, Jimmy Lin, and Dina Demner-Fushman. 2006. Evaluation of pico as a knowledge representation for clinical questions. AMIA Annual Symposium Proceedings, 2006:359–363.

Qiao Jin, Bhuwan Dhingra, Zhengping Liu, William Cohen, and Xinghua Lu. 2019a. PubMedQA: A dataset for biomedical research question answering. In Proceedings of the 2019 Conference on Empirical Methods in Natural Language Processing and the 9th International Joint Conference on Natural Language Processing (EMNLP-IJCNLP), pages 2567–2577, Hong Kong, China. Association for Computational Linguistics.

Qiao Jin, Bhuwan Dhingra, Zhengping Liu, William W. Cohen, and Xinghua Lu. 2019b. Pubmedqa: A dataset for biomedical research question answering. In Proceedings of the 2019 Conference on Empirical Methods in Natural Language Processing and the 9th International Joint Conference on Natural Language Processing, EMNLP-IJCNLP 2019, Hong Kong, China, November 3-7, 2019, pages 2567–2577. Association for Computational Linguistics.

Martin Krallinger, Obdulia Rabal, Saber A Akhondi, Martin Pérez Pérez, Jesús Santamaría, Gael Pérez Rodríguez, Georgios Tsatsaronis, Ander Intxaurrondo, José Antonio López, Umesh Nandal, et al. 2017. Overview of the biocreative vi chemical-protein interaction track. In Proceedings of the sixth BioCreative challenge evaluation workshop, volume 1, pages 141–146.

Jiao Li, Yueping Sun, Robin J. Johnson, Daniela Sciaky, Chih-Hsuan Wei, Robert Leaman, Allan Peter Davis, Carolyn J. Mattingly, Thomas C. Wiegers, and Zhiyong Lu. 2016. Biocreative V CDR task corpus: a resource for chemical disease relation extraction. Database J. Biol. Databases Curation, 2016.

Pengfei Liu, Weizhe Yuan, Jinlan Fu, Zhengbao Jiang, Hiroaki Hayashi, and Graham Neubig. 2023. Pretrain, prompt, and predict: A systematic survey of prompting methods in natural language processing. ACM Computing Surveys, 55(9):1–35.

Anastasios Nentidis, Konstantinos Bougiatiotis, Anastasia Krithara, and Georgios Paliouras. 2020a. Results of the seventh edition of the bioasq challenge. In Machine Learning and Knowledge Discovery in Databases: International Workshops of ECML PKDD 2019, Würzburg, Germany, September 16–20, 2019, Proceedings, Part II, pages 553–568. Springer.

Anastasios Nentidis, Konstantinos Bougiatiotis, Anastasia Krithara, and Georgios Paliouras. 2020b. Results of the seventh edition of the bioasq challenge. CoRR, abs/2006.09174.

Benjamin Nye, Junyi Jessy Li, Roma Patel, Yinfei Yang, Iain J Marshall, Ani Nenkova, and Byron C Wallace. 2018. A corpus with multi-level annotations of patients, interventions and outcomes to support language processing for medical literature. In Proceedings of the conference. Association for Computational Linguistics. Meeting, volume 2018, page 197. NIH Public Access.

Long Ouyang, Jeffrey Wu, Xu Jiang, Diogo Almeida, Carroll Wainwright, Pamela Mishkin, Chong Zhang, Sandhini Agarwal, Katarina Slama, Alex Ray, et al. 2022. Training language models to follow instructions with human feedback. Advances in Neural Information Processing Systems, 35:27730–27744.

Yifan Peng, Shankai Yan, and Zhiyong Lu. 2019. Transfer learning in biomedical natural language processing: An evaluation of BERT and elmo on ten benchmarking datasets. In Proceedings of the 18th BioNLP Workshop and Shared Task, BioNLP@ACL 2019, Florence, Italy, August 1, 2019, pages 58–65. Association for Computational Linguistics.

Long N Phan, James T Anibal, Hieu Tran, Shaurya Chanana, Erol Bahadroglu, Alec Peltekian, and Gré-goire Altan-Bonnet. 2021. Scifive: a text-to-text transformer model for biomedical literature. arXiv preprint arXiv:2106.03598.

Alec Radford, Karthik Narasimhan, Tim Salimans, Ilya Sutskever, et al. 2018. Improving language understanding by generative pre-training.

Alec Radford, Jeffrey Wu, Rewon Child, David Luan, Dario Amodei, Ilya Sutskever, et al. 2019. Language models are unsupervised multitask learners. OpenAI blog, 1(8):9.

Colin Raffel, Noam Shazeer, Adam Roberts, Katherine Lee, Sharan Narang, Michael Matena, Yanqi Zhou, Wei Li, and Peter J Liu. 2020. Exploring the limits of transfer learning with a unified text-to-text transformer. The Journal of Machine Learning Research, 21(1):5485–5551.

Kalhan Rosenblatt. 2023. Chatgpt passes mba exam given by a wharton professor. Retrieved Jan, 25:2023.

Larry Smith, Lorraine K Tanabe, Cheng-Ju Kuo, I Chung, Chun-Nan Hsu, Yu-Shi Lin, Roman Klinger, Christoph M Friedrich, Kuzman Ganchev, Manabu Torii, et al. 2008. Overview of biocreative ii gene mention recognition. Genome biology, 9(2):1–19.

Gizem Soğancioğlu, Hakime Öztürk, and Arzucan Özgür. 2017. Biosses: a semantic sentence similarity estimation system for the biomedical domain. Bioinformatics, 33(14):i49–i58.

Jason Wei, Xuezhi Wang, Dale Schuurmans, Maarten Bosma, Ed Chi, Quoc Le, and Denny Zhou. 2022. Chain of thought prompting elicits reasoning in large language models. arXiv preprint arXiv:2201.11903.

Michihiro Yasunaga, Jure Leskovec, and Percy Liang. 2022. Linkbert: Pretraining language models with document links. In Proceedings of the 60th Annual Meeting of the Association for Computational Linguistics (Volume 1: Long Papers), ACL 2022, Dublin, Ireland, May 22-27, 2022, pages 8003–8016. Association for Computational Linguistics.

